# Host-derived population genomics data provides insights into bacterial and diatom composition of the killer whale skin

**DOI:** 10.1101/282038

**Authors:** Rebecca Hooper, Jaelle C. Brealey, Tom van der Valk, Antton Alberdi, John W. Durban, Holly Fearnbach, Kelly M. Robertson, Robin W. Baird, M. Bradley Hanson, Paul Wade, M. Thomas, P. Gilbert, Phillip A. Morin, Jochen B.W. Wolf, Andrew D. Foote, Katerina Guschanski

## Abstract

Recent exploration into the interactions and relationship between hosts and their microbiota has revealed a connection between many aspects of the host’s biology, health and associated microorganisms. Whereas amplicon sequencing has traditionally been used to characterise the microbiome, the increasing number of published population genomics datasets offer an underexploited opportunity to study microbial profiles from the host shotgun sequencing data. Here, we use sequence data originally generated from killer whale *Orcinus orca* skin biopsies for population genomics, to characterise the skin microbiome and investigate how host social and geographic factors influence the microbial community composition. Having identified 845 microbial taxa from 2.4 million reads that did not map to the killer whale reference genome, we found that both ecotypic and geographic factors influence community composition of killer whale skin microbiomes. Furthermore, we uncovered key taxa that drive the microbiome community composition and showed that they are embedded in unique networks, one of which is tentatively linked to diatom presence and poor skin condition. Community composition differed between Antarctic killer whales with and without diatom coverage, suggesting that the previously reported episodic migrations of Antarctic killer whales to warmer waters associated with skin turnover may control the effects of potentially pathogenic bacteria such as *Tenacibaculum dicentrarchi*. Our work demonstrates the feasibility of microbiome studies from host shotgun sequencing data and highlights the importance of metagenomics in understanding the relationship between host and microbial ecology.

## 1 INTRODUCTION

The skin microbiome is an ecosystem comprised of trillions of microbes sculpted by ecological and evolutionary forces acting on both the microbes and their host (McFall-Ngai, & Eliot, 2005; Byrd et al., 2018). Recent explorations have revealed a tight connection between almost every aspect of the host’s biology and the associated microbial community (Reviewed by McFall-Ngai, & Eliot, 2005; Bordenstein, & Theis, 2015; Alberdi et al., 2016; Koskella et al., 2017; Byrd et al., 2018). Although numerous intrinsic and extrinsic factors that influence the skin microbiome composition have been identified, the relative importance of these factors often appear to differ even between closely related host taxa (Kueneman et al., 2014; McKenzie, Bowers, Fierer, Knight, & Lauber, 2012; Wolz et al., 2017). Intrinsically, the host’s evolutionary history, age, sex, and health appear significant (Leyden, McGiley, Mills, & Kligman, 1975; Cho & Blaser, 2012; McKenzie et al., 2012; Phillips et al., 2012; Apprill et al., 2014; Ying et al., 2015; Chng et al., 2016). Extrinsically, both environmental factors, where a sub-selection of environmental microbes colonise host skin (Apprill et al., 2014; Wolz et al., 2017; Walke et al., 2014; Ying et al., 2015), and socioecological factors, such as a host’s social group and the level of interaction with conspecifics (Jain, 2011; Lax et al., 2014; Song et al., 2013; Kolodny et al. 2017), can play important roles.

Most microbiome studies to date are based on 16S ribosomal RNA gene sequences, a highly conserved region of the bacterial and archaeal genome (Hamady & Knight, 2009). However, in addition to potential biases in PCR amplification, in which low reliability of quantitative estimations arise due mismatches in primer binding sites, PCR stochasticity, and different numbers of 16S gene copies in each bacterial species (Alberdi et al., 2018), analysis of the 16S region can limit functional and taxonomic classification (Quince, Walker, Simpson, Loman, & Segata, 2017). In contrast, shotgun metagenomics can facilitate both high-resolution taxonomic and functional analyses (Ranjan, Rani, Metwally, McGee, & Perkins, 2016; Quince, Walker, Simpson, Loman, & Segata, 2017; Koskella et al., 2017). The advent of affordable high-throughput sequencing has seen an ever-increasing number of population genomics studies in a wide range of study systems (e.g. Jones et al., 2012; Poelstra et al., 2014; Der Sarkissian et al., 2015; Nater et al., 2017). This affords an unprecedented opportunity to exploit sequencing data to secondarily investigate the microbial communities associated with the sampled tissue of their host (Salzberg et al., 2005; Ames et al., 2015; Zhang et al., 2015; Mangul et al., 2016; Lassalle et al., 2018).

Here, we explore the relative importance of extrinsic factors on the epidermal skin microbiome of free-ranging killer whales (*Orcinus orca*) using shotgun sequencing data derived from skin biopsy samples of five ecologically specialised populations, or ecotypes (Foote et al., 2016). Given the widespread geographic range (Forney & Wade, 2007) and variation in ecological specialisation of killer whales, even in sympatry (Durban et al., 2017; Ford et al., 1998), this species provides a good study system for exploring the effects of both geographic location and ecotype (a proxy for both sociality and phylogenetic history) on the skin microbiome. However, the opportunistic use of such data is also fraught with potential pitfalls. We therefore describe in detail measures taken to disentangle potential sources of contamination from the true skin microbiome, thus providing a useful roadmap for future host-microbiome studies that exploit host-derived shotgun sequencing data.

## 2 MATERIALS AND METHODS

### 2.1 Study system

Throughout most of the coastal waters of the North Pacific two ecotypes of killer whales are found in sympatry: the mammal-eating *‘transient’* and fish-eating *‘resident’* ecotype (Ford et al., 1998; Saulitis et al., 2000; Matkin et al., 2007; Filatova et al., 2015). Four decades of field studies have found that they are socially and genetically isolated (Hoelzel & Dover, 1991; Barrett-Lennard, 2000; Hoelzel et al., 2007; Ford, 2009; Parsons et al., 2013; Filatova et al., 2015; Morin et al., 2015; Foote & Morin, 2016). Killer whales have also diversified into several ecotypes in the waters around the Antarctic continent, including a form commonly observed hunting seals in the pack-ice of the Antarctic peninsula (*type B1*), a form that feeds on penguins in the coastal waters of the Antarctic peninsula (*type B2*), and a dwarf form thought to primarily feed on fish in the dense pack-ice of the Ross Sea (*type C*) (Pitman & Ensor, 2003; Pitman & Durban, 2010, 2012; Durban et al., 2017; Pitman et al. 2018).

### 2.2 Sample collection and data generation

We used the unmapped reads from a published population genomics study of killer whale ecotypes (European Nucleotide Archive, www.ebi.ac.uk/ena, accession numbers: ERS554424-ERS554471; Foote et al. 2016), which produced low coverage genomes from a total of 49 wild killer whales, corresponding to five ecotypes; 10 samples each of the North Pacific fish-eating *resident* and sympatric mammal-eating *transient* ecotypes, and 8, 11 and 10 samples, respectively from Antarctic types *B1, B2* and *C* (see Figure 1 for the sampling locations). DNA was extracted from epidermal biopsies collected by firing a lightweight dart with a sterilised stainless-steel cutting tip from a sterilised projector (e.g. Palsbøll et al., 1991; Barrett-Lennard et al., 1996) at the flank of the killer whale. As a study on captive killer whales found low variability in the taxonomic composition of the skin microbiome from different body sites (Chiarello et al. 2017), small variation in the exact location on the flank from which the biopsy was taken should not bias our results. Biopsies were stored in sterile tubes at −20°C. At no point were biopsy samples in direct contact with human skin.

**FIGURE 1.**
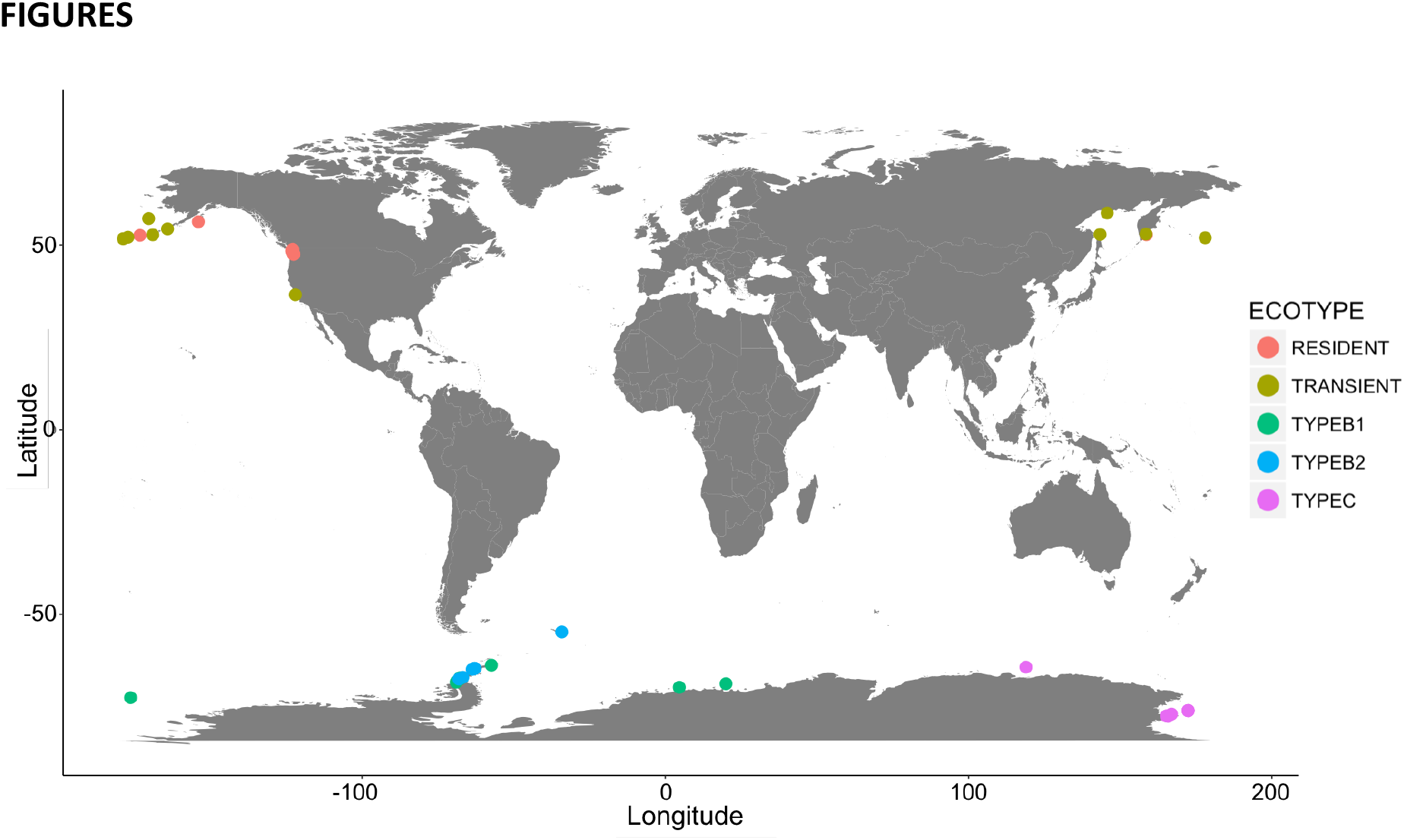
Summary of geographic sample origins and microbial diversity measures. Map of sampling locations of the 49 samples of five killer whale ecotypes, from which skin microbiomes were included in this study. The Antarctic ecotypes primarily inhabit waters 8-16°C colder than the North Pacific ecotypes.

DNA extraction, library building, and sequencing have been previously described (Foote et al., 2016). All lab work was conducted in a sterile flow hood to prevent contamination. Sequencing was done at the Danish National High-Throughput DNA Sequencing Centre within the University of Copenhagen. The facility is specifically geared for low DNA quantity library sequencing from ancient and environmental DNA. Samples of the same ecotype were pooled and sequenced across multiple sequencing lanes. Samples of different ecotypes were always run on different sequencing lanes, with the exception of several *type B1* and *B2* samples, which were initially grouped as *‘type B’* (Pitman & Ensor, 2003) and some samples were therefore sequenced on shared lanes.

### 2.3 Sequencing read pre-processing

As a means to enrich the dataset for bacterial sequences, we first used SAMtools v1.5 (Li et al., 2009) to remove all sequencing reads that mapped to the killer whale reference nuclear genome (Oorca1.1, GenBank: ANOL00000000.2; Foote et al., 2015) and mitochondrial genome (GU187176.1) with BWA-mem (Li & Durbin, 2009). The remaining reads were adapter-trimmed using AdapterRemoval V2.1.7 (Schubert, Lindgreen, & Orlando, 2016). We then removed duplicates generated during library indexing PCR by merging reads with identical sequences using in-house python scripts (Dryad doi:10.5061/dryad.c8v3rv6). All reads with an average quality score <30 were filtered out using prinseq V0.20.4 (Schmieder & Edwards, 2011), and all reads of <35bp were removed using AdapterRemoval.

### 2.4 Investigating contamination

Despite the precautions outlined above, contamination can be introduced at several stages of the sequence data generation, and subsequently mistaken for the genuine host-associated microbiome signal. Contaminating DNA can be present in laboratory reagents and extraction kits (Lusk et al., 2014; Salter et al., 2014). For example, silica in some commercial DNA spin columns is derived from diatom cells and therefore can be a potential source of contamination with diatom DNA (Naccache et al., 2013). However, the Qiagen QIAquick spin columns used in this study do not contain silica from biological material, according to the manufacturer. Cross contamination can also occur between samples processed in the same sequencing centre (Ballenghien et al., 2017). The impact of contamination increases in samples with small amounts of true exogenous DNA and can swamp the signal from the host’s microbiome (Lusk et al., 2014; Salter et al., 2014). Contamination can be assessed using negative controls (e.g. Davis et al., 2017). However, the data used in this study was initially produced with the sole focus on the host organism. Including extractions and library preparation blanks is not a routine procedure in population genomics studies based on high-quality host tissue samples and, as such, blanks were not included in the laboratory workflow and hence not sequenced. Therefore, we instead implement an *ad hoc* workflow that attempts to differentiate between contaminant and real exogenous DNA from host species shotgun sequencing data.

#### 2.4.1 PhiX contamination

The contamination of microbial reference genomes by PhiX, which is used as a control in Illumina sequencing, is a known potential source of error in metagenomics studies using shotgun sequencing data (Mukherjee et al., 2015). Therefore, to avoid erroneous mapping of PhiX-derived reads to contaminated genomes, we removed all reads mapping to the PhiX genome used by Illumina (NC_001422) with BWA-mem 0.7.15 (Li, 2013) with default parameters.

#### 2.4.2 Environmental and laboratory contamination

If the amount of contamination (derived from laboratory reagents or environment) is relatively equal among samples, we expect the relative proportion of contaminant sequencing reads to be inversely correlated to the quantity of sample-derived DNA: *i.e*. low DNA quantity samples will be disproportionately affected by contaminant DNA sequences compared with high-quantity samples (Lusk et al., 2014; Salter et al., 2014). We therefore estimated the correlation between the proportion of the total sequencing reads assigned to each microbial taxon (see below for how taxonomic assignment was conducted) and total DNA read count per sample (prior to removal of host DNA and before PCR duplicate removal). Microbial taxa for which the read count was significantly negatively correlated with the total number of reads per sample (including host DNA), *i.e*. those that consistently increased in abundance in low-quantity DNA samples, were flagged as potential contaminants.

#### 2.4.3 Human contamination

To account for the possibility of contamination with human-associated microorganisms, we next quantified the amounts of human DNA in our samples and used this as a proxy of human-derived microbial contamination (see Supporting Information for the details of read processing). Only reads uniquely mapping to a single region of the genome with high quality (samtools -q 30 -F 4 -F 256) were retained, and we removed all duplicates using samtools rmdup in single-end mode. Human contamination levels were estimated by calculating the percentage of filtered reads mapped to the human genome (Table S1). We included these values as a covariate in statistical models as a way to, at least partially, control for contamination with human-associated microorganisms.

#### 2.4.4 Known bacterial contaminants

Next, we investigated whether specific bacterial taxa that have previously been reported to be likely contaminants are present in our dataset. Following read-based analyses, we found that our samples were dominated by *Cutibacterium (Propionibacterium) acnes*, which is abundant on human skin (Byrd et al., 2018) and a known contaminant of high-throughput sequencing data (Lusk et al., 2014; Mollerup et al., 2016). We therefore investigated the distribution of sequence identity between our *C. acnes* reads and the *C. acnes* reference genomes, with the expectation that human or laboratory contaminants would show high (close to 100) percent identity, whereas killer whale-derived *C. acnes* would be more divergent.

Additionally, we analysed data from a North Pacific killer whale sequenced at ~20x coverage in a published study, in which sample collection, DNA extraction, and sequencing was entirely independent of our data production (accession number: SRP035610; Moura et al., 2014). If *C. acnes* was present in these data, it would suggest that either it was a real component of the killer whale skin microbiome, or it was independently introduced as contamination in both studies.

Contaminant taxa are unlikely to be introduced in isolation. C. acnes was confirmed to be a likely contaminant (see below) and we therefore removed all taxa with which it significantly co-occurred. Using netassoc 0.6.3 (Morueta-Holme et al., 2016), we calculated co-occurrence scores between all taxon pairs in the raw taxa dataset. We set the number of null replicates to 999, and corrected p-values for multiple comparisons using the FDR method. From the resulting matrix, we selected taxa with the top 10% absolute significant co-occurrence score with candidate contaminant taxa, and removed these taxa from downstream analyses, along with *C. acnes*.

#### 2.4.5 Investigating sources of contamination

Finally, to ascertain the authenticity of our data and to estimate the level and possible source of contamination, we used SourceTracker V2.0.1 (Knights et al., 2011), a tool that implements a Bayesian classification model to predict the proportion of taxa derived from different potential source environments. This approach allowed us to compare the composition of the free-ranging killer whale skin microbiome to other marine mammal skin microbiota and to a number of potential contaminating and environmental sources. We obtained data from public repositories and included microbial communities reflecting the marine environment (ocean water from Southern Ocean and the North Pacific, Sunagawa et al., 2015), other marine mammal skin (captive bottlenose dolphins *Tursiops truncatus* and killer whales along with the respective pool water samples, and free-ranging humpback whales, Chiarello et al., 2017, Bierlich et al., 2018), likely contaminants such as human skin and gut (Oh et al., 2014, Lloyd-Price et al., 2017, Meisel et al., 2016), and laboratory contamination from commonly used reagents (sterile water, Salter et al., 2014) (Table S2). We attempted to specifically select sources that were obtained with the shotgun sequencing approach to avoid potential locus-specific effects that can produce distinct microbiome profiles in amplicon-based studies. However, only 16S rRNA amplicon data was available for the marine mammal skin and the laboratory contaminants, each study targeting a different region within this locus (Table S2). Therefore, to control for locus-specific effects, we also included samples from a human skin 16S amplicon study (Meisel et al., 2016) and limited our data to reads mapping to the 16S rRNA gene for those comparisons (see Supporting Information for more detailed methodology of read processing).

We used the R package Vegan V2.4.6 (Oksanen et al., 2017) to calculate distances between microbiome profiles derived from these different datasets. After Total Sum Scaling (TSS) normalization, abundance-based Bray-Curtis and presence/absence-based binary Jaccard distances were calculated and visualised using principal coordinates analysis. Subsequently, a subset of sources was used in SourceTracker and we used our killer whale data as sink without applying rarefaction to either sink or source samples. We also repeated the SourceTracker analysis using free-ranging humpback whales as the sink samples.

### 2.5 Taxonomic assignment

We used MALT (MEGAN Alignment Tool) version 0.3.8 (Herbig et al., 2016) to create a reference database of bacterial genomes downloaded from the NCBI FTP server (ftp://ftp.ncbi.nlm.nih.gov/genomes/all/GCA, accessed 26^th^ January 2017). We performed a semi-global nucleotide-nucleotide alignment against the reference database. Semi-global alignments are more suitable for assessing quality and authenticity criteria common to short-read data, and are also useful when aligning 16S rRNA data against a reference database such as SILVA (Herbig et al., 2016). Sequence identity threshold was set to 95% as per Vågene et al., (2018), but with a more conservative threshold of including only taxa with five or more aligned reads in subsequent analysis,

The nucleotide alignments produced in MALT were further analysed in MEGAN version 6.7.6 (Huson et al., 2016). Genomes with the presence of stacked reads in some genomic regions and/or large gaps without any mapped reads, were flagged using a custom python script (Dryad doi:10.5061/dryad.c8v3rv6), and manually assessed in MEGAN. This step was necessary to identify spurious and incorrectly supported bacterial taxa, which were removed from further analysis if they represented highly abundant species (Warinner et al., 2017). Taxonomic composition of the samples was also interactively explored in MEGAN, and the number of reads assigned to each taxon was exported for subsequent analysis.

Taxonomic assignment was also carried out using an assembly-based approach. Filtered metagenomic sequences of all samples were merged to perform a co-assembly using Megahit 1.1.1 (Li et al., 2015) with default settings and k-list: 21,29,39,59,79. Assembly quality was assessed using Quast 4.5 (Gurevich et al., 2013). Contigs were subsequently mapped to reference bacterial genomes with MGMapper (Petersen et al., 2017) using best mode to assign taxonomy. The assembly file was indexed using BWA-index and Samtools-faidx. BWA-mem was subsequently used to map the reads of each sample back to the assembly contigs to finally retrieve the mapped reads using SAMtools-view. Individual coverage values were calculated with Bedtools 2.26.0 (Quinlan et al., 2010) and contig coverage table normalised using Cumulative Sum Scale (CSS) as implemented in MetagenomeSeq (Paulson et al., 2013). The sequencing data used is this study is rather shallow in terms of coverage of microbial taxa, corresponding to low coverage killer whale genomes (mean of 2x). Therefore, we explored how low sequencing depth may affect the inferred bacterial profiles. To this end, we used an independently sequenced 20x coverage *resident* killer whale genome (Moura et al., 2014). By drawing a random subset of reads from this genome using SAMtools, we compared the taxonomic composition of the microbiome of the same individual at 20x, 10x, 5x and 2x mean sequence coverage depth.

### 2.6 Diversity analyses

We calculated all diversity measures in Vegan (Oksanen et al., 2017), using reads that were assigned to the species level in MEGAN. By focussing on taxa at the species-level, we were able to explore the skin microbiome at a high resolution, an advantage of shotgun over amplicon-based analyses. However, results of this analysis should be interpreted in light of a species-level focus, where we are exploring a small yet well-resolved representation of the microbiome, which may potentially be enriched with pathogens and common environmental bacteria, rather than a holistic representation of the entire ecosystem.

To control for bias introduced by varying genome size (species with larger genomes show higher read counts, which are translated into higher abundance scores; Warinner et al., 2017), we divided all read counts by the size of the respective full bacterial genome. If the taxon was mapped to the level of the strain, we divided the read count by the published genome size of that strain; if identified to the species level, we divided the read count by the average genome size across all published strains of that species.

Beta diversity was explored using two dissimilarity matrices in the R-package Vegan: abundance-based Bray-Curtis and presence/absence-based binary Jaccard distances. To assess the strength and significance of ecotype and geographic location (longitude and latitude) in describing variation in community composition, we conducted permutational multivariate analysis using the function *Adonis* in Vegan. We controlled for differing depths of coverage between samples using two techniques. First, we used genome size-controlled data (see above) and included the number of reads mapping to the species level as a covariate. Second, TSS normalisation of the genome-size controlled data was conducted, followed by conversion to the Bray-Curtis distance matrix. TSS normalisation is irrelevant for presence absence data, as only species presence, rather than species abundance, is retained in the binary presence absence matrix. As a result, three models were explored: two Bray-Curtis models with differing depth-control techniques, and one Jaccard model using read counts as covariate. Each model consisted of the following covariates: latitude (numeric), longitude (numeric), ecotype (categorical), and percentage human contamination (numeric), with library size included only when TSS normalisation was not used. For each model, residuals were permuted 9999 times. We used the function *Betadisper* (Vegan) followed by ANalysis Of VAriance (ANOVA) to test for homogeneity of group dispersions. *Betadisper* can be used to ensure that i) the *Adonis* model results are not confounded by heterogeneous variances (Anderson, 2001) and ii) to make biological inferences about between-group variance in community composition.

We used the function *capscale* from the Vegan package to perform Principal Co-ordinate Analysis (PCoA). The four bacterial taxa that described the most variation on PCoA1, and the four that described the most variation on PCoA2, were designated as ‘driving taxa’. We therefore classified a total of eight unique driving taxa that describe individual differences in microbiome composition (Table S4).

### 2.7 Network analysis

To venture beyond single microbial taxa and explore microbial interactions that include interspecific dynamics, we expanded our analyses to networks of bacterial communities associated with the driving taxa identified through the PCoA. Using netassoc (Morueta-Holme et al., 2016), we compared the observed partial correlation coefficients between taxa with a null distribution estimated from identical species richness and abundances as the observed data. Again, taxa co-occurrence scores were calculated between all taxon pairs in the raw dataset, with null replicates set to 999. The FDR method was used to correct p-values for multiple comparisons. From the resulting matrix of significant co-occurrence scores, we selected the 20 taxa with the highest absolute co-occurrence score for each of the 8 unique driving taxa. We created a new matrix including only these taxa and visualised co-occurrence networks.

### 2.8 Functional profiling

Community composition can be a poor predictor of the functional traits of the microbiome, due to processes such as horizontal gene transfer (HGT) between bacterial taxa, which can decouple species composition and function (Koskella et al., 2017). Shifting focus from the taxonomic composition to the genic composition of the microbiome reduces the impact of HGT on functional characterisation (Koskella et al., 2017).

To explore functional profiles of the samples, we used DIAMOND v0.9.10 with default parameters (Buchfink, Xie & Huson, 2015) to create a reference database of non-redundant protein sequences from fully sequenced bacterial genomes downloaded from the NBCI FTP server (https://ftp.ncbi.nlm.nih.gov/genomes/genbank/ accessed 9th March 2017). Nucleotide-to-amino-acid alignments of the sample reads to the reference database was performed in DIAMOND and the top 10% of alignments per query reported. The MEGAN tool daa-meganizer was then used to assign reads to proteins based on the DIAMOND alignments and to assign functional roles to these proteins using the SEED (Overbeek et al., 2005) and eggNOG (Huerta-Cepas et al., 2017) databases. Since one protein can have more than one function, it is possible for one read to be assigned to multiple functional subsystems. The raw count data (number of reads assigned to functional subsystem) were exported from MEGAN and further processed in R. To control for differences in library depth, read counts per functional group were normalised by total read numbers mapping to SEED or eggNOG terms. We used principal components analysis (PCA) performed in the R package *prcomp* to visualise differences in functional groups between individuals.

We additionally performed an assembly-based functional profiling to overcome the individual weaknesses of both assembly and read-based methodologies (Quince et al., 2017). *Ab initio* gene prediction was performed over the metagenomic assembly using Prodigal 2.6.3 (Hyatt et al., 2010). The list of predicted gene sequences was indexed using BWA and SAMtools was used to map the reads of each sample back to the gene sequences. We used Bedtools 2.26.0 (Quinlan et al., 2010) to calculate individual coverages. Gene coverage table was subsequently CSS normalised using MetagenomeSeq (Paulson et al., 2013).

### 2.9 Diatom-association analyses

Antarctic killer whales are often observed to have a yellow hue, which has been attributed to diatom coverage (Berzin & Vladimirov, 1983; Pitman & Ensor, 2003) and identifiable individuals have been observed to transit from this yellow skin colouration to a ‘clean’ skin condition (Durban & Pitman, 2012). This change is hypothesised to occur during brief migrations to subtropical latitudes, where turnover of the outer skin layer takes place with a reduced thermal cost (Durban & Pitman, 2012). If this hypothesis is correct, diatom abundance should be correlated to skin age and colouration (Hart, 1935; Konishi et al. 2008; Durban & Pitman, 2012). Inter-individual variation in microbiome profiles within the Antarctic ecotypes could therefore reflect variation in the age of the outer skin layer. During network analysis, we identified a possible association between a key bacterial taxa driving between-sample differences in community composition (*Tenacibaculum dicentrarchi*) and bacterial taxa associated with diatoms. Following from our observations that three samples from Antarctic ecotypes had high abundances of *T. dicentrarchi* and that in the PCoA these samples were differentiated from most other samples, we investigated the link between observed diatom coverage, abundance of *T. dicentrarchi* and abundance of other algae-associated bacterial taxa. We conducted qualitative colour grading of *type B1* and *type B2* individuals using photographs taken at the time of biopsy collection, ranging from ‘clean’ through to ‘prominent’ yellow colouration.

Only a limited number of diatom reference genomes are available and therefore we used two methodologies to quantify the level of diatom DNA in our samples. First, we used MALT and MEGAN in the same taxonomic pipeline as previously described, but with a reference database comprised of NCBI RefSeq nucleotide sequences from the diatom phylum Bacillariophyta (downloaded 30th October 2017). To date, only seven diatom reference genomes are available, thus identification at the species level was not attempted. Instead, numbers of reads mapping to Bacillariophyta were exported and further processed in R. Raw diatom read counts were converted to proportion of total number of sequencing reads per sample. Second, we aligned all reads against the SILVA rRNA database (release 128, Quast et al., 2013) using BWA-mem 0.7.17, retained reads mapping with > 10 mapping quality with SAMtools and used uclust (Edgar, 2010) in QIIME 1.9.1 (Caporaso et al., 2010) to assign taxonomy based on the SILVA 18S database at 97% similarity. From the resulting OTU table we retained reads that matched to known diatom taxonomy.

We explored the correlation between latitude (by grouping North Pacific and Antarctic ecotypes) and proportion of reads per sample mapping to diatoms using a generalised linear model with a quasipoisson error structure and log link. As covariates, we included longitude, number of reads mapping to the bacterial species level to control for library size, and number of human reads to control for human-associated microbial contamination. Using the same model structure, we then tested the correlation between proportion of reads per sample mapping to diatoms and the presence/abundance of *T. dicentrarchi* reads, as well as the correlation with presence of known algae-associated bacterial taxa (including *T. dicentrarchi, Cellulophaga baltica, Formosa* sp. *Hel1_33_131, Winogradskyella* sp., *Marinovum algicola, Agarivorans gilvus, Pseudoalteromonas atlantica* and *Shewanella baltica:* Bowman 2000; Amin et al., 2012; Goecke et al., 2013a,b, incorporated as a binary variable).

## 3 RESULTS

Metagenomic profiles from the skin microbiome of 49 killer whales from five ecotypes (Figure 1) were successfully reconstructed using shotgun sequencing data from DNA extracted from skin biopsies. Of the reads retained following our stringent filtering procedure, but before our investigations into *C. acnes* as a possible contaminant, 8.20% (n = 7,984,195) were assigned to microbial taxa using the read-based approach, with 2.45% (n = 2,384,587) assigned at the species level (see Dryad repository doi:10.5061/dryad.c8v3rv6). Overall, 845 taxa of microbes were identified. The co-assembly yielded a 33.01 Mbp-long metagenome comprised of 45,934 contigs (N50 = 970 bp, average = 730 bp, max = 48,182 bp). Taxonomy was assigned to 41.73% of the contigs. Results from the assembly-based approach were concordant with the read-based results, we therefore report only the latter.

### 3.1 Investigating contamination

On average 0.16% of reads (range 0.01-5.43%) reads mapped to the human genome (Table S2), suggesting presence of human contamination and making it possible that human-derived bacteria were present in our dataset. After correcting for multiple testing, we found no significant negative correlation between the proportion of reads assigned to each bacterial taxon and the total number of sequenced reads (Supporting Information, Fig. S1). Negative trends (although not significant) between some bacterial taxa and the total number of sequenced reads were largely driven by one outlier sample with the lowest coverage (B1_124047). Following the deduplication step of our processing pipeline, these taxa were no longer present in the dataset, as they fell below our defined threshold of five aligned reads in MALT (Fig. S2).

*C. acnes* was identified as the most abundant bacterial taxon, with an average abundance of 39.57% (standard deviation = 24.65; Fig. S3), but it may have been introduced via human or laboratory contamination (Lusk et al., 2014). Percent identity to the human-derived *C. acnes* genome was 100% for 245 and over 97% for 505 of the 527 contigs identified as *C. acnes* by MGMapper (Fig. S4), supporting the idea of a likely exogenous source of *C. acnes*. Killer whale samples pooled by ecotype were sequenced across multiple sequencing lanes, allowing us to investigate if contamination with *C. acnes* was introduced at the sequencing step. Relative *C. acnes* abundance per sample was highly similar between sequencing lanes (Coefficient of Variation = 0.076; Fig. S5), suggesting that the contamination occurred prior to sequencing. However, *C. acnes* was also present to a high abundance (18.06% of reads aligning at species level) in the independently sequenced *resident* killer whale (Moura et al., 2014), suggesting that contamination with *C. acnes* was not specific to our workflow. We concluded that there was a high probability that *C. acnes* was a lab contaminant and therefore removed all *C. acnes* reads/contigs from our dataset before continuing with analysis.

#### 3.1.1 Network analysis results for C. acnes associated taxa

Following its identification as a likely contaminant, we used network analysis to identify and remove the top 10% of species which significantly co-occurred with *C. acnes*, which corresponded to co-occurrence scores above the absolute value of 1000 (Fig. S6). Overall, 82 species were removed (Dryad doi:10.5061/dryad.c8v3rv6), many of which are known human-associated bacterial taxa. Following this filtering step, one *type C* sample had no remaining taxa. We therefore excluded this sample from further analyses.

#### 3.1.2 Metagenomic affinities of wild killer whale skin microbiome

Only 10 killer whale samples had 50 or more 16S reads with assigned SILVA taxonomy (eight killer whale samples remained after filtering for *C. acnes* associated taxa, Figure 2). Overall, prior to C. acnes filtering, the killer whale dataset had 273 taxa in common with the dataset of 2,279 bacterial taxa derived from sources (e.g. human, marine mammal and environmental samples, see section 2.4.5). After filtering for *C. acnes* and associated taxa, 236 of the 273 killer whale associated taxa remained. Free-ranging killer whale and humpback whale skin microbiomes overlapped on the principal coordinates, independent of the applied distance measure and presence of *C. acnes* associated bacteria (Figure 2a, Fig. S7a,b). In contrast, data from the captive study, including killer whale and captive dolphin skin samples and their pool water, clustered separately from all other studies. General separation by sequencing approach (i.e. shotgun versus amplicon) was not observed: For instance, amplicon- and shotgun-sequenced human samples grouped together (Figure 2a,b). It is therefore possible that the separation of the captive study samples is due to either the use of a specific 16S target locus or other factors associated with captive versus wild environments (note that the pool was filled with sea water from the Mediterranean Sea; Chiarello et al., 2017).

**Figure 2.**
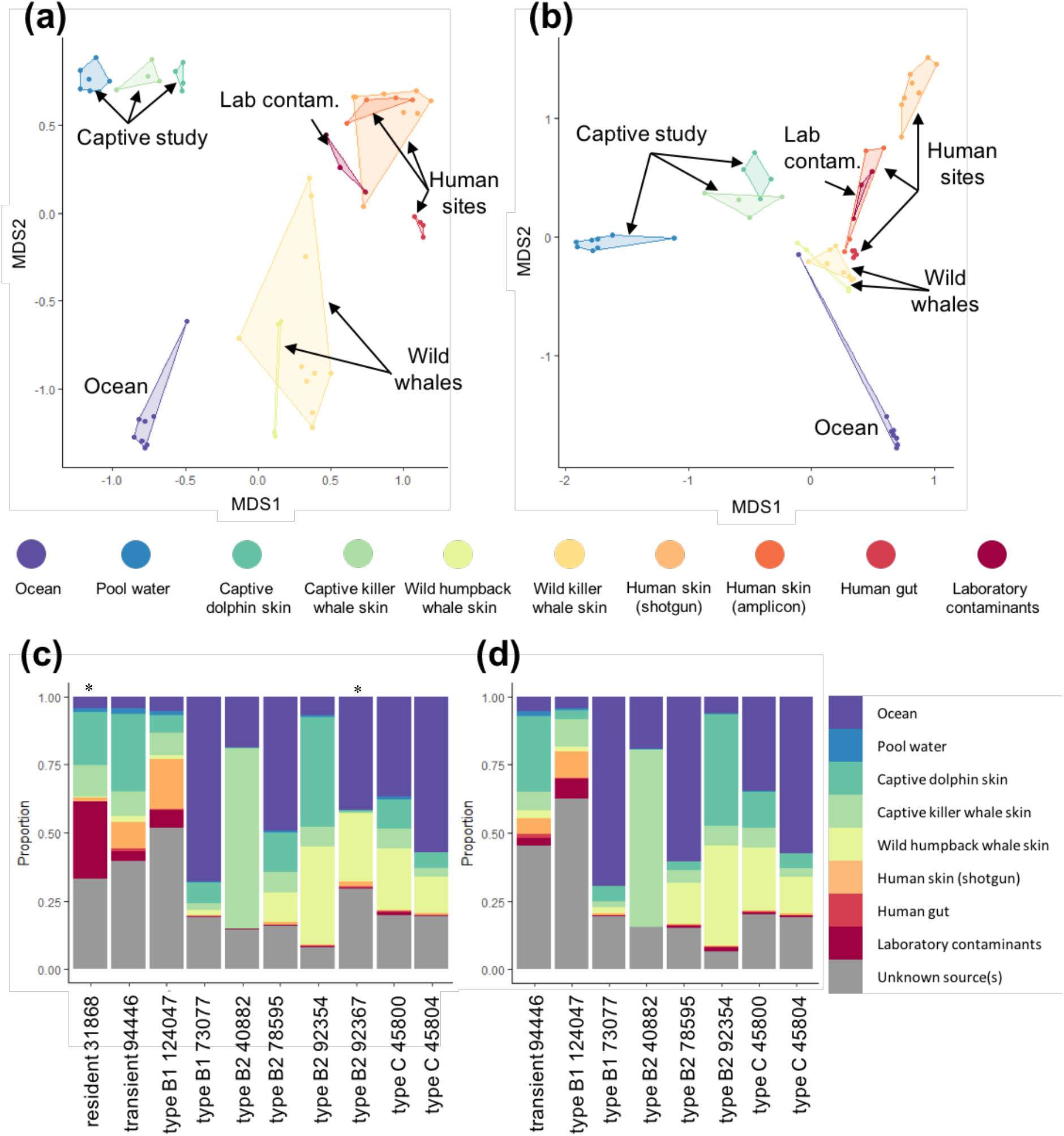
Composition of the wild killer whale skin microbiomes and other published microbiomes, for samples with ≥ 50 taxonomy assigned 16S reads. Principal coordinates analysis of Jaccard binary presence/absence distances before **(a)** and after **(b)** filtering of *C. acnes* associated taxa from the wild killer whale data. Proportions of sources contributing to each killer whale sample, represented by columns, from SourceTracker analysis before **(c)** and after **(d)** filtering of *C. acnes* associated taxa. * in **(c)** denotes samples that were excluded after *C. acnes* filtering due to low read numbers.

The three marine mammal species formed one cluster irrespective of study on the third dimension in the abundance-based Bray-Curtis distance analysis (Fig. S7c,d), suggesting that there is a common factor to the marine mammal skin microbiome composition. Importantly, the free-ranging killer whale microbiome profiles generally grouped away from the human skin samples, gut samples and laboratory contaminants. They were also separated from the ocean water samples, suggesting that the killer whale skin microbiomes characterised in our study represent a microbial community that is clearly distinct from surrounding ocean water. Here, it is noteworthy that filtering of our data for *C. acnes* associated taxa at the genus level is highly conservative and also removes a number of microbial taxa that are abundant in the marine environment, as they belong to the same genera as some *C. acnes* associated species. Samples representing laboratory contamination consistently clustered with the human skin samples (Figure 2a,b, Fig. S7), suggesting that one source of contaminants in laboratory work are human-associated skin microbes. All results presented above were confirmed with a larger dataset that included 16 killer whale samples with at least 20 bacterial 16S reads with SILVA taxonomy assignment (Fig. S8).

Based on the principal coordinates analysis and for greater clarity of presentation, we restricted the selection of samples that were used as sources in the SourceTracker analysis to captive dolphin skin (n=4), captive killer whale skin (n=4), water from the captive killer whale pool (n=4), wild humpback whale skin (n=4), Southern Ocean water (n=4), human gut (n=4), shotgun-derived human skin data from a sebaceous site (n=4), and laboratory contamination (n=3; the fourth sample had < 20 16S reads and was excluded from analysis) (Table S2). The SourceTracker results supported those of the principal coordinates analysis (Figure 2c,d), with human skin taxa contributing on average only 3.4% to the wild killer whale skin microbiome (range 0.0-18.4%). This percentage decreased to 2.2% (range 0.0-9.6%) after filtering out *C. acnes* associated taxa. The contribution of lab contaminants was also low (average 4.2%, range 0.0-28.6) in all but one resident killer whale individual (31868), which was removed after *C. acnes* filtering due to low (< 50) read numbers (average 1.7%, range 0.0-7.1% after removal of *C. acnes* associated taxa). The sources contributing the most to the free-ranging killer whale skin microbiomes after removing *C. acnes* associated taxa included Southern Ocean (mean 32.3%, range 4.5-69.4%), humpback whale skin (11.9%, range 0-36.7% in), captive killer whale skin and captive dolphin skin (mean 13.2%, range 2.1-64.8% and mean 12.5%, range 0.2-40.8%, respectively). A high proportion of taxa observed in free-ranging killer whales could not be assigned to any of the sources included in the analysis (‘Unknown’, mean >25%). These taxa may represent uncharacterised diversity specific to the wild killer whale skin microbiome, a source that was not included in our analysis, e.g. ocean water collected at the same time as the killer whale skin biopsies, or marine mammal skin taxa that are poorly characterised by the 16S locus targeted in other marine mammal microbiome studies.

To verify the SourceTracker results for free-ranging killer whale samples studied here, we also ran SourceTracker using the four wild humpback whales as the sink samples while assigning free-ranging killer whales as a source (Fig. S9). Two humpback whales sampled early in the foraging season around the Antarctic Peninsula closely resembled the wild killer whale profiles, containing a mixture of taxa attributed to the wild killer whale skin (41.7% and 65.3%), the captive dolphin skin (31.1% and 2.7%) and unknown sources (21.3% and 24.5%). In contrast, the microbiome of the two humpback whales sampled late in the Antarctic foraging season were dominated by Southern Ocean taxa (both >95%). This is consistent with the temporal variation in the complete humpback whale dataset reported by Bierlich et al., (2018). Overall, the detailed analyses of contributing sources of the killer whale skin microbiome revealed a large proportion of taxa that are also found on the skin of other marine mammals and an important contribution of environmental ocean water taxa. This is in line with previous reports that found a significant contribution of sea water to, yet distinct composition of, marine mammal microbiomes (Bik et al., 2016). Expected contaminating sources, such as human skin and laboratory contaminants, contributed only a small proportion to our killer whale skin microbiome data obtained from host shotgun sequencing.

### 3.2 Taxonomic exploration

Read-based and assembly-based approaches produced concordant taxonomic profiles. The most abundant constituents of the killer whale skin microbiome at the phylum level were Proteobacteria, Actinobacteria, Bacteroidetes and Firmicutes (Fig. S3a), which have been identified in previous studies of baleen whale skin microbiota (Appril et al., 2014; Shotts et al., 1990), including through 16S amplification of skin swabs from captive killer whales under controlled conditions (Chiarello et al., 2017). At the species level, we found a high level of inter-individual variation (Figure 3a, Fig. S3b), as previously found for four captive killer whales housed in the same facility (Chiarello et al., 2017).

**FIGURE 3.**
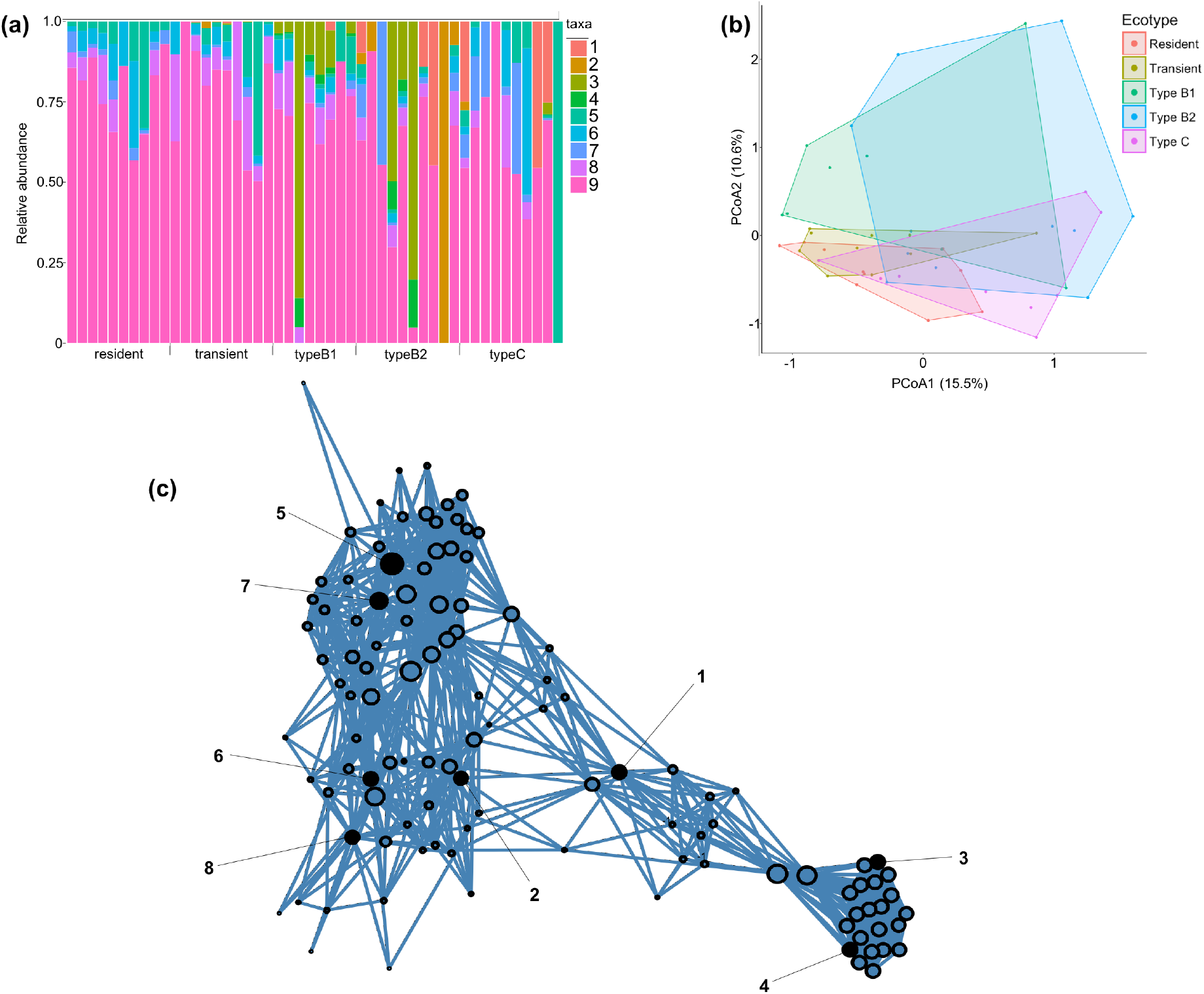
**(a)** Proportion of driving bacteria per individual, after data filtering. Individuals, represented by columns, are grouped by ecotype, and the relative proportions of bacterial taxa are indicated by column shading (1, *Tenacibaculum dicentrarchi;* 2, *Paraburkholderia fungorum*; 3, *Pseudoalteromonas haloplanktis*; 4, *Pseudoalteromonas translucida*; 5, *Acinetobacter johnsonii*; 6, *Pseudomonas stutzeri*; 7, *Stenotrophomonas maltophilia*; 8, *Kocuria palustris*; 9, Other). **(b)** Beta diversity between ecotypes illustrated as a Bray-Curtis PCoA estimated from read counts. **(c)** Positive co-occurrence network built from a cooccurrence matrix of all species, subsetted to the 8 driving taxa (black nodes numbered as above) and their top 20 positive and significant co-occurring species. Only species with a significant co-occurrence score of >800 are shown.

Subsetting an independently sequenced *resident* killer whale genome to lower sequencing depth, we inferred that while five most common taxa were found in similar proportions in high and low coverage data, the identification of rarer taxa became more stochastic at lower sequencing depths (Table S3). Our results may therefore suffer from this bias associated with low coverage data, which would be most prominent in presence/absence-based analyses. As a means to control for this bias, we include library size as a covariate in models investigating beta diversity.

### 3.3 Diversity analyses

Human contamination was not a significant driver in the models exploring beta diversity (Table 1), explaining at most 2% of the variation in taxonomic composition in each model. Ecotype was a significant variable in all models, explaining 10-11% of variation in the data (Table 1). Latitude was significant in both Bray Curtis models but not in the Jaccard presence-absence model. Where significant, it explained 4 – 5% of variation in the data (Table 1). Longitude was not significant in any of the models. *Betadisper* analysis revealed no significant heterogeneity in the variation of community composition between ecotypes (non-TSS normalised Bray-Curtis: df = 4, F = 0.52, p = 0.72; TSS normalised Bray-Curtis: df = 4, F = 1.74, p = 0.16; binary Jaccard: df = 4, F = 0.63, p = 0.64). This suggests that between-individual variation in microbial composition is similar among ecotypes.

**Table 1.** Factors influencing the killer whale skin microbiome. Results of Adonis models using genome-size controlled species data. (a) Bray Curtis model with library size included as a covariate; (b) TSS normalised Bray Curtis model; (c) Binary Jaccard model with library size included as a covariate. Significant factors are highlighted in bold.

|  | (a) Bray Curtis |  |  | (b) Bray Curtis<br>(TSS normalised) |  |  | (c) Binary Jaccard |  |  |
| --- | --- | --- | --- | --- | --- | --- | --- | --- | --- |
|  | *F* | *r*<sup>2</sup> | *p* | *F* | *r*<sup>2</sup> | *p* | *F* | *r*<sup>2</sup> | *p* |
| Latitude | 1.8 | 0.04 | **0.01** | 2.35 | 0.05 | **<0.01** | 1.4 | 0.03 | 0.05 |
| Longitude | 0.89 | 0.02 | 0.62 | 0.89 | 0.02 | 0.61 | 0.99 | 0.02 | 0.45 |
| Ecotype | 1.36 | 0.11 | **<0.01** | 1.35 | 0.11 | **0.02** | 1.23 | 0.10 | **0.03** |
| Library size | 1.33 | 0.03 | **<0.01** | - | - | - | 1.20 | 0.02 | 0.23 |
| Human contamination | 1.13 | 0.02 | 0.35 | 0.59 | 0.01 | 0.89 | 1.04 | 0.02 | 0.42 |
| Residuals |  | 0.79 |  |  | 0.81 |  |  | 0.80 |  |
| Total |  | 1 |  |  | 1 |  |  | 1 |  |

The Bray-Curtis PCoA explained more variation than Jaccard (24.13% versus 16.06% on the first two axes), we therefore focus on the Bray-Curtis results. A network based on significant co-occurrences between eight bacterial taxa driving variation at the individual level (Table S4) and the top 20 co-occurring taxa for each of the driving taxa, showed clearly differentiated and distinct community groups (Figure 3). Further investigation into the top 20 taxa that significantly co-occurred with the driving taxa found that three of the taxa showing the highest co-occurrence scores with the driving taxon *T. dicentrarchi (Formosa sp. Hel1_33_131, Cellulophaga algicola* and *Algibacter alginolytica*) are associated with algae (Bowman, 2000; Becker et al., 2017; Sun et al., 2016).

### 3.4 *T. dicentrarchi* and diatoms

Both approaches to diatom identification produced concordant results (Fig. S10, Table S5). Antarctic ecotypes had a significantly higher abundance of diatom DNA than North Pacific ecotypes (β = 0.65, SE = 0.29, p = 0.03; Figure 4a). Individuals with “prominent” yellow colouration showed higher diatom abundance (Figure 4b), supporting the link between skin colour and diatom presence in Antarctic killer whales. Furthermore, the abundance of diatom DNA per sample was significantly positively correlated with the abundance and presence of *T. dicentrarchi* reads (number of *T. dicentrarchi* reads: β = 0.014, SE = 0.003, p = <0.001; presence of *T. dicentrarchi:* β = 0.915, SE = 0.207, p <0.001; Figure 4c) and the presence of at least one algae-associated bacterial taxa (β = 0.98, SE = 0.17, p = <0.001; Figure 4d).

**FIGURE 4.**
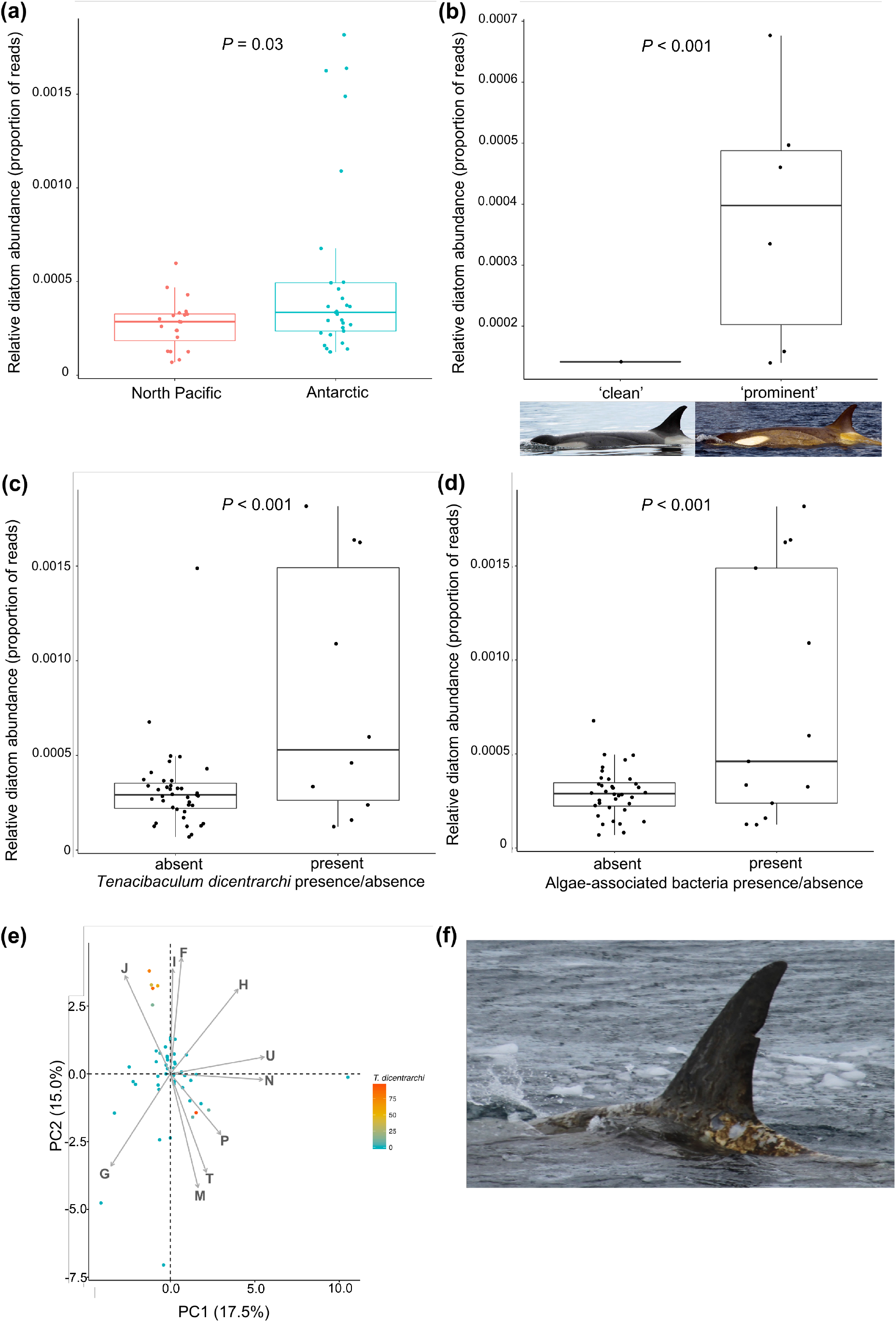
The influence of diatom abundance on skin microbiome community composition and microbial functional profiles. **(a)** Relative diatom abundance is significantly higher in Antarctic killer whales than North Pacific, but this is largely driven by a subset of outlier Antarctic individuals. **(b)** Within Antarctic *type B1* and *type B2* specimens, the relative diatom abundance is significantly associated with skin colouration of the host killer whale, with the yellowish hue being a reliable indicator of diatom load. Inset images are of the same *type B2* killer whale individual displaying extreme variability in diatom coverage, both photographs by John Durban. Relative diatom abundance is significantly associated with **(c)** the presence of *Tenacibaculum dicentrarchi* and **(d)** several algae-associated bacteria, including *T. dicentrarchi*. **(e)** PCA of variation in functional COGs between individuals, coloured by *T. dicentrarchi* abundance. Individuals with high relative abundances of *T. dicentrarchi* generally cluster with high values in principal component 2. The top 10 COGs contributing to PCA variation are shown in grey arrows (J: translation, ribosomal structure and biogenesis, I: lipid transport and metabolism, F: nucleotide transport and metabolism, H: coenzyme transport and metabolism, U: intracellular trafficking, secretion and vesicular transport, N: cell motility, P: inorganic ion transport and metabolism, T: signal transduction mechanisms, M: cell wall membrane envelope biogenesis, G: carbohydrate transport and metabolism). **(f)** Photograph of a *type B1* killer whale in the Gerlache Strait of the Antarctic Peninsula on the 4th December 2015 with high diatom coverage and poor skin condition. Photograph by Conor Ryan.

### 3.5 Functional analysis

In the read-based functional analysis a total of 3,611,441 reads mapped to eggNOG functions and 1,440,371 reads mapped to SEED functions. In the contig-based functional analysis we identified 56,042 potential genes in our metagenome, out of which EggNog function was assigned to 35,182. Both approaches identified energy production and conversion (class C) and amino acid metabolism and transport (class E) as the most abundant eggNOG functions in our dataset. The eggNOG PCA revealed high variability between individuals (Figure 4e), however, a cluster of Antarctic whales was observed in principal component 2. These samples had high abundances of *T. dicentrarchi* (Figure 4e) and were associated with functions corresponding to the COG functional categories J (translation, ribosomal structure and biogenesis, β = 0.007, SE = 0.002, p = <0.001), F (nucleotide transport and metabolism, β = 0.005, SE = 0.002, p = 0.004) and I (lipid transport and metabolism, β = 0.004, SE = 0.002, p = 0.03). The same cluster of high abundance *T. dicentrarchi* Antarctic samples was also identified in the SEED PCA (Fig. S11). These samples had increased numbers of reads mapping to DNA metabolism, amino acids and derivatives and cofactors/vitamins, although none of these functions were significantly correlated with *T. dicentrarchi* abundance.

## 4 DISCUSSION

Our study highlights that communities of exogenous or host-associated microbiota can be genetically characterised from shotgun sequencing of DNA extracted from the host tissue. However, dedicated analysis and treatment of contamination is necessary and requires careful consideration in studies such as this, whereby samples were not collected nor sequenced with the intention of genetically identifying microbiota. In such cases, the normal stringent control measures which are routine in microbial studies, such as the sequencing of blanks, may not be possible. We have therefore presented an array of approaches for estimating the proportion and sources of contamination and accounting for it in shotgun studies. Overall, our analyses suggest that with careful consideration, the mining of microbial DNA from host shotgun sequencing data can provide useful biological insights that inform future targeted investigations into microbiome composition and function under stringent laboratory conditions.

After carefully filtering our data, we were able to identify species interactions, ecological networks and community assembly of the microbes and diatoms that colonise killer whale skin by utilising unmapped reads from shotgun sequencing data generated from skin biopsies. A key advantage of this approach over amplicon-based sequencing is the ability to assess functional variation based on gene content, and to identify taxa to species level (Koskella et al., 2017; Quince, Walker, Simpson, Loman, & Segata, 2017). However, despite ongoing efforts to describe bacterial species diversity, the breadth of the reference database is a limiting factor in the unbiased characterisation of bacterial composition. Thus, taxa identified in our analyses are necessarily limited to species with available genomic information, and in some cases are likely to represent their close phylogenetic relatives (Tessler et al., 2017). Hence, we refer to ‘taxa’ rather than ‘species’ where appropriate. We also demonstrate the impact of contamination on the low numbers of reads from true host-associated microbes, which can dilute the signal of biologically meaningful variation among samples.

Social and geographic factors have been found to influence microbial diversity in terrestrial and semi-terrestrial animals (Koskella et al., 2017). However, there is less understanding of how these factors interplay in a wide-ranging social marine mammalian system (Nelson et al., 2015). We found that beta diversity of the killer whale skin microbiome was significantly influenced by ecotype and latitude. Temperature has been shown to be a key determinant of marine microbial community structure at a global scale (Salazar & Sunagawa, 2017; Sunagawa et al., 2015). However, the effect of ecotype as the most important tested variable highlights the significance of social and phylogenetic factors in shaping microbiome richness and composition. In addition, it underscores that although killer whale skin is influenced by the local environment (Romano-Bertrand et al., 2015), it represents a unique ecosystem that is separate from that of the surrounding habitat. Concordant with our results, a study of the microbiome of four captive killer whales and the sea water from their pool found that the skin microbiota were more diversity and phylogenetically distinct from the sea water microbial community (Chiarello et al., 2017). Killer whales are highly social mammals (Baird, 2000; Ford, 2009), and thus are likely to have a high potential for horizontal transfer of microbes between individuals during contact (Nelson et al., 2015). Ecotype-specific social behaviour, organisation and population structure, as well as other variables related to ecotype ecology, such as range size, and diet (due to transmission of bacteria from different prey species; Menke et al., 2017), are all likely to affect the diversity of microbial species that individuals are exposed to, and also influence the level of horizontal transfer of microbes between whales. The strong social philopatry in killer whales (Baird, 2000; Ford, 2009) and the phylogenetic and phylogeographic history of ecotypes is also likely to play a role, whereby due to limited social transmission between ecotypes, the phylogeny of bacterial species is likely to reflect that of the host (Ley et al., 2008, but see Rothschild et al., 2018). It is also likely to be influenced by the host’s evolutionary history, including secondary contact between ecotypes (Foote & Morin, 2016), where both vertical and horizontal transmission of microbes between ecotypes is possible.

Despite the significance of ‘ecotype’ as a driver of skin microbiome diversity in killer whales, at least 79% of the variation in the microbiome is unexplained by the drivers considered in our models (Table 1). There is a strong overlap among ecotypes in the PCoA (Figure 3b), suggesting a shared core microbiome which may be partially shared with other cetacean species (Figure 2). Additionally, the PCoA shows substantial variation within ecotypes (Figure 3b), further highlighting the role of some other driver(s) of microbiome variation. Among Antarctic ecotypes, individual variation was associated with diatom presence and a discrete subnetwork of microbial taxa. The occurrence of a ‘yellow slime’ attributed to diatoms on the skin of whales, including killer whales, was recorded as early as a century ago (Bennett 1920; Pitman et al., 2018). The extent of diatom adhesion on Antarctic whales is thought to correlate with latitude and the time the whale has spent in cold waters (Hart, 1935; Konishi et al. 2008). The skin microbiome of humpback whales has been reported to change through the Antarctic foraging season (Bierlich et al., 2018), and our SourceTracker analysis found that humpback whales sampled during the late foraging season (*i.e*. individuals who had presumably spent longer in the Southern Ocean waters at the time of sampling) had more similarity to Southern Ocean microbial communities than those collected during the early foraging season. This raises the intriguing question as to whether the time spent in the frigid Antarctic waters could be a driver of variation in the skin microbiome and diatom load of Antarctic killer whales.

Satellite tracking of *type B1* and *B2* killer whale movements, Durban & Pitman (2012) documented rapid return migrations to subtropical latitudes, in which individuals travelled up to 9,400 km in 42 days. Based on the strong directionality and velocity of travel during these migrations, Durban & Pitman (2012) hypothesised that they were not associated with breeding or feeding behaviour. Instead, they argued that these migrations could be driven by the need to leave the frigid Antarctic waters and temporarily move to warmer waters, to allow for physiological maintenance including the regeneration of the outer skin layer (Durban & Pitman, 2012). The identification of the same individuals in Antarctic waters, sometimes with a thick accumulation of diatoms, and at other times appearing ‘clean’, supports the hypothesis that skin regeneration is an intermittent rather than continuous process (Durban & Pitman, 2012). Furthermore, bite marks from cookiecutter sharks *Isistius spp.*, which are typically found in warm and temperate waters, are regularly observed on *type C* killer whales (Pitman et al., 2018), which have also been directly tracked to warm sub-tropical waters (Durban et al., 2013).

We present genetic support for the hypothesis of Durban & Pitman (2012), that ‘clean’ and yellow-tinted *type B1* and *B2* killer whales represent differences in diatom load. In addition, we provide the first evidence that the extent of diatom coverage is also associated with significant variation in the skin microbiome community. We found that Antarctic killer whales with the highest diatom abundance also had skin microbiomes most similar to Southern Ocean microbial communities, suggesting that at the time of sampling these individuals had spent longer in the Antarctic waters, consistent with the hypothesis that diatom coverage accumulates with time spent in the cold Southern Ocean waters. Perhaps most significantly, diatom abundance was positively correlated with the abundance of T. *dicentrarchi*, a known pathogen in several fish species, which is associated with skin lesions and severe tail and fin rot (Piñeiro-Vidal et al., 2012; Habib et al., 2014; Avendaño-Herrera et al., 2016).

Our analyses revealed that samples with high abundances of T. *dicentrarchi* show distinct functional profiles. Functional analyses remain exploratory at this stage, constrained by the difficulty of translating broad functional categories into biological meaning. However, with more data that links individual health status and microbiome composition, functional analyses may provide a tool for identifying individuals at risk. Therefore, whether T. *dicentrarchi* represents a pathogen to killer whale hosts remains unknown. Type *B1* killer whales in apparently poor health and with heavy diatom loads have been observed with severe skin conditions (skin peeling and lesions; Figure 4f), however, *Tenacibaculum* sp. have been reported in up to 95% of humpback whales sampled in recent studies, which included apparently healthy individuals (Appril et al., 2011, 2014; Bierlich et al., 2018). Skin maintenance may thus represent a balancing act for Antarctic killer whales of managing the costs of pathogen load, thermal regulation, reduced foraging time and long-range movement. Research into the skin microbiome should therefore continue to form a component of the ongoing holistic and multidisciplinary research programme to investigate the health of Antarctic killer whale populations and more broadly in studies on the health of marine mammals (e.g. Appril et al., 2014; Raverty et al., 2017).

Ongoing field efforts provide the opportunity to further explore the relationships and interactions between killer whale hosts, their skin microbiome, other exogenous symbionts such as diatoms, and the environment. Our community-based analyses suggest the presence of a distinct environmental taxa network centred on *P. haloplanktis* as a driving taxon (Fig. 3c). Collection and metagenomic characterization of environmental samples, such as sea water, alongside host biological samples would allow further explorations into the contribution of local ecological factors to the host microbiome. As a means of reducing the impact of contamination with DNA from laboratory environment, microbiome characterisation can be conducted by means of RNA sequencing. This has an additional advantage of generating metatranscriptomic data, which, in combination with the metagenomic data, can facilitate the comparison/contrast between community function (using RNA transcript) and community taxonomic composition (using DNA sequence; Koskella et al., 2017). This may further reduce the potential impact of common lab contaminants, allowing the exploration of the bacterial functional repertoire that is in use in a given ecological context, including reconstruction of metabolic pathways (Bashiardes, Zilberman-Schapira, Elinav 2016). Contamination in the laboratory could be further controlled for and characterised through inclusion of extraction, library preparation and PCR blanks as negative controls (Lusk et al., 2014; Salter et al., 2014) and measures such as double-indexing (Rohland & Reich, 2012; van der Valk et al., 2018), which can then inform the emerging downstream filtering methods for separating true microbiomes from contamination (Delmont & Eren, 2016; Davis et al., 2018). Lastly, the advances in long-read sequencing using portable nanopore-based platforms make it possible to generate data suitable for reconstructing complete bacterial genomes while in the field (Parker et al., 2017), including in the Antarctic (Johnson et al., 2017). This is a promising development with respect to improving the breadth of host taxa from which bacterial taxa are derived and should improve future mapping of metagenomics data and taxonomic assignment.

## Acknowledgements

The suggestion of harvesting the skin microbiome from host shotgun data was first mooted by Gerald Pao of the Salk Institute during a meeting back in 2012 when we first embarking on the shotgun sequencing project, and we are grateful to Gerald for sowing this seed. We would like to thank Bob Pitman who was involved in the collection of many of the samples used in this study and greatly contributed through many discussions on the variation among killer whale types. We’d further like to thank David Studholme for pointing out the rich literature surrounding diatom microbiomes, as well as potential diatom-related contamination. James Fellows Yates, Linda Rhodes, Morten Limborg and the microbiome journal club of the Evogenomics section at the Centre for GeoGenetics provided valuable feedback on this work and an earlier draft of this manuscript. We acknowledge The Danish National High-Throughput DNA Sequencing Centre for sequencing the samples and, particularly, Andaine Seguin-Orlando, Lillian Petersen, Cecilie Demring Mortensen, Kim Magnussen and Ian Lissimore for technical support. Sample collection from killer whales in Antarctica was supported by the Lindblad Expeditions-National Geographic Conservation Fund. This work was funded by European Research Council grant ERCStG-336536 to J.B.W.W.; a Danish National Research Foundation grant DNRF94 to M.T.P.G, the Welsh Government and Higher Education Funding Council for Wales through the Sêr Cymru National Research Network for Low Carbon, Energy and Environment, and from the European Union’s Horizon 2020 research and innovation programme under the Marie Skłodowska-Curie grant agreement No. 663830 and a British Ecological Society grant (SR17\1227) to AF, the FORMAS grant (project 2016-00835) to KG. A.A. was supported by the Danish Council for Independent Research - DFF (grant 5051-00033) and Lundbeckfonden (grant R250-2017-1351). Finally, we would like to thank the Linnaeus Scholarship Foundation at the University of Uppsala for providing a scholarship to RH, the Erasmus Mundus Master Programme in Evolutionary Biology (MEME) for facilitating the collaborative supervision of RH, and to Vanessa and Peter Hooper for providing financial support to RH throughout this project.

## Data Accessibility

All sequencing data are archived at the European Nucleotide Archive, www.ebi.ac.uk/ena, accession numbers: ERS554424-ERS554471. Scripts and metadata are available in Dryad repository doi:10.5061/dryad.c8v3rv6.

## Author Contributions

R.H., J.B., T.V. and A.A. analysed the data; J.W.D. and H.F. conducted the photographic grading. A.F. and K.G. conceived and coordinated the study, which was developed from a suggestion by Gerald Pao of the Salk Institute. J.W.D., H.F., R.W.B., M.B.H. and P.W. were involved in sample collection and DNA was extracted by K.M.R. R.H., J.B., A.F., K.G. wrote the manuscript with input from T.V., A.A., J.W.D., H.F., R.W.B., M.T.P.G., P.A.M. and J.B.W.W.

## References

Alberdi, A., Aizpurua, O., Bohmann, K., Zepeda-Mendoza, M. L., & Gilbert, M. T. P. (2016). Do vertebrate gut metagenomes confer rapid ecological adaptation?. Trends in ecology & evolution, 31, 689–699.

Alberdi, A., Aizpurua, O., Gilbert, M. T. P., & Bohmann, K. (2017). Scrutinizing key steps for reliable metabarcoding of environmental samples. Methods in Ecology and Evolution, 9, 134–147.

Ames S. K., Gardner S. N., Marti J. M., Slezak T. R., Gokhale M. B., Allen J. E. (2015) Using populations of human and microbial genomes for organism detection in metagenomes. Genome Research, 25, 1056–1067.

Amin, S. A., Parker, M. S., & Armbrust, E. V. (2012). Interactions between diatoms and bacteria. Microbiology and Molecular Biology Reviews, 76(3), 667–684.

Anderson, M. J. (2001). A new method for non-parametric multivariate analysis of variance. Austral ecology, 26(1), 32–46.

Apprill, A., Robbins, J., Eren, A. M., Pack, A. A., Reveillaud, J., Mattila, D., Moore, M., Niemeyer, M., Moore, K.M. & Mincer, T. J. (2014). Humpback whale populations share a core skin bacterial community: towards a health index for marine mammals?. PLoS One, 9(3), e90785.

Avendaño-Herrera, R., Irgang, R., Sandoval, C., Moreno-Lira, P., Houel, A., Duchaud, E., … & Ilardi, P. (2016). Isolation, characterization and virulence potential of Tenacibaculum dicentrarchi in salmonid cultures in Chile. Transboundary and emerging diseases, 63(2), 121–126.

Baird, R. W. (2000). The killer whale. Cetacean societies: field studies of dolphins and whales, 127–153.

Ballenghien, M., Faivre, N., & Galtier, N. (2017). Patterns of cross-contamination in a multispecies population genomic project: detection, quantification, impact, and solutions. BMC biology, 15(1), 25.

Barrett-Lennard, L. G. (2000). Population structure and mating patterns of killer whales (Orcinus orca) as revealed by DNA analysis (Doctoral dissertation, University of British Columbia).

Barrett-Lennard, L., Smith, T. G., & Ellis, G. M. (1996). A cetacean biopsy system using lightweight pneumatic darts, and its effect on the behavior of killer whales. Marine Mammal Science, 12(1), 14–27.

Bashiardes S, Zilberman-Schapira G, Elinav E. (2016). Use of Metatranscriptomics in Microbiome Research. Bioinformatics and Biology Insights, 10, 19–25.

Becker, S., Scheffel, A., Polz, M. F., & Hehemann, J. H. (2017). Accurate quantification of laminarin in marine organic matter with enzymes from marine microbes. Applied and environmental microbiology, 83(9), e03389–16.

Belden, L. K., Hughey, M. C., Rebollar, E. A., Umile, T. P., Loftus, S. C., Burzynski, E. A., Minbiole, K. P., House, L. L., Jensen, R. V., Becker, M. H. & Walke, J. B. (2015). Panamanian frog species host unique skin bacterial communities. Frontiers in microbiology, 6, 1171.

Bennett, A. G. (1920). On the occurrence of diatoms on the skin of whales. In Proc. R. Soc. Lond. B (Vol. 91, No. 641, pp. 352–357).

Berzin, A. A., & Vladimirov, V. L. (1983). A new species of killer whale (Cetacea, Delphinidae) from the Antarctic waters. Zoologichesky Zhurnal, 62(2), 287–295.

Bik, E. M., Costello, E. K., Switzer, A. D., Callahan, B. J., Holmes, S. P., Wells, R. S., … Relman, D. A. (2016). Marine mammals harbor unique microbiotas shaped by and yet distinct from the sea. Nature Communications, 7, 10516. doi:10.1038/ncomms10516

Bista, I., Carvalho, G. R., Tang, M., Walsh, K., Zhou, X., Hajibabaei, M., … & Christmas M. (2018). Performance of amplicon and shotgun sequencing for accurate biomass estimation in invertebrate community samples. Molecular ecology resources.

Bolger AM, Lohse M, Usadel B (2014) Trimmomatic: a flexible trimmer for Illumina sequence data. Bioinformatics, 30, 2114–20.

Bordenstein, S. R., & Theis, K. R. (2015). Host biology in light of the microbiome: ten principles of holobionts and hologenomes. PLoS biology, 13(8), e1002226.

Bowman, J. P. (2000) Description of Cellulophaga algicola sp. nov., isolated from the surface of Antarctic algae, and reclassification of Cytophaga uliginosa (ZoBell and Upham 1944) Reichenbach 1989 as Cellulophaga uliginosa comb. nov. International Journal of Systematic and Evolutionary Microbiology, 50, 1861–1868.

Buchfink, B., Xie, C., & Huson, D. H. (2015). Fast and sensitive protein alignment using DIAMOND. Nature methods, 12(1), 59.

Buller, N. B. (2014). Bacteria and fungi from fish and other aquatic animals: a practical identification manual. Cabi.

Byrd, A. L., Belkaid, Y., & Segre, J. A. (2018). The human skin microbiome. Nature Reviews Microbiology.

Caporaso, J. G., Kuczynski, J., Stombaugh, J., Bittinger, K., Bushman, F. D., Costello, E. K., … & Huttley, G. A. (2010). QIIME allows analysis of high-throughput community sequencing data. Nature methods, 7(5), 335.

Chng, K. R., Tay, A. S. L., Li, C., Ng, A. H. Q., Wang, J., Suri, B. K., Matta, S. A., McGovern, N., Janela, B., Wong, X. F. C. C. & Sio, Y. Y. (2016). Whole metagenome profiling reveals skin microbiome-dependent susceptibility to atopic dermatitis flare. Nature microbiology, 1(9), 16106.

Csardi, G., & Nepusz, T. (2006). The igraph software package for complex network research. InterJournal, Complex Systems, 1695(5), 1–9.

Davis, N. M., Proctor, D., Holmes, S. P., Relman, D. A., & Callahan, B. J. (2017). Simple statistical identification and removal of contaminant sequences in marker-gene and metagenomics data. bioRxiv, 221499.

Delmont, T. O., & Eren, A. M. (2016). Identifying contamination with advanced visualization and analysis practices: metagenomic approaches for eukaryotic genome assemblies. PeerJ, 4, e1839.

Durban, J. W., Fearnbach, H., Burrows, D. G., Ylitalo, G. M., & Pitman, R. L. (2017). Morphological and ecological evidence for two sympatric forms of Type B killer whale around the Antarctic Peninsula. Polar Biology, 40(1), 231–236.

Durban, J. W., & Pitman, R. L. (2012). Antarctic killer whales make rapid, round-trip movements to subtropical waters: evidence for physiological maintenance migrations?. Biology Letters, 8(2), 274–277.

Durban JW, Pitman RL (2013) Out of Antarctica: dive data support ‘physiological maintenance migration’ in Antarctic killer whales. 20th Biennial Conference on the Biology of Marine Mammals. Society for Marine Mammalogy (9-13 December 2013, Dunedin, New Zealand).

Edgar, R. C. (2010). Search and clustering orders of magnitude faster than BLAST. Bioinformatics, 26(19), 2460–2461.

Filatova, O. A., Borisova, E. A., Shpak, O. V., Meshchersky, I. G., Tiunov, A. V., Goncharov, A. A., Fedutin, I. D. & Burdin, A. M. (2015). Reproductively isolated ecotypes of killer whales *Orcinus orca* in seas of the Russian far east. Biology Bulletin of the Russian Academy of Sciences, 42, 674–681.

Foote, A. D., & Morin, P. A. (2016). Genome-wide SNP data suggest complex ancestry of sympatric North Pacific killer whale ecotypes. Heredity, 117(5), 316.

Foote, A. D., Vijay, N., Ávila-Arcos, M. C., Baird, R. W., Durban, J. W., Fumagalli, M., … & Robertson, K. M. (2016). Genome-culture coevolution promotes rapid divergence of killer whale ecotypes. Nature communications, 7, 11693.

Ford, J. K., Ellis, G. M., Barrett-Lennard, L. G., Morton, A. B., Palm, R. S., & Balcomb III, K. C. (1998). Dietary specialization in two sympatric populations of killer whales (Orcinus orca) in coastal British Columbia and adjacent waters. Canadian Journal of Zoology, 76(8), 1456–1471.

Ford, J. K. (2009). Killer whale: Orcinus orca. In Encyclopedia of Marine Mammals (Second Edition) (pp. 650–657).

Forney, K. A., & Wade, P. R. (2006). Worldwide distribution and abundance of killer whales. Whales, whaling and ocean ecosystems, 145–162.

Goecke, F., Labes, A., Wiese, J., & Imhoff, J. F. (2013a). Phylogenetic analysis and antibiotic activity of bacteria isolated from the surface of two co-occurring macroalgae from the Baltic Sea. European journal of phycology, 48(1), 47–60.

Goecke, F., Thiel, V., Wiese, J., Labes, A., & Imhoff, J. F. (2013b). Algae as an important environment for bacteria-phylogenetic relationships among new bacterial species isolated from algae. Phycologia, 52(1), 14–24.

Gurevich, A., Saveliev, V., Vyahhi, N., & Tesler, G. (2013). QUAST: quality assessment tool for genome assemblies. Bioinformatics, 29(8), 1072–1075.

Habib, C., Houel, A., Lunazzi, A., Bernardet, J. F., Olsen, A. B., Nilsen, H., … & Duchaud, E. (2014). Multilocus sequence analysis of the marine bacterial genus *Tenacibaculum* suggests parallel evolution of fish pathogenicity and endemic colonization of aquaculture systems. Applied and environmental microbiology, 80(17), 5503–5514.

Hamady, M., & Knight, R. (2009). Microbial community profiling for human microbiome projects: Tools, techniques, and challenges. Genome research, 19(7), 1141–1152.

Hart, T. J. (1935). On the diatoms of the skin film of whales, and their possible bearing on problems of whale movements. University Press.

Herbig, A., Maixner, F., Bos, K. I., Zink, A., Krause, J., & Huson, D. H. (2016). MALT: Fast alignment and analysis of metagenomic DNA sequence data applied to the Tyrolean Iceman. bioRxiv, 050559.

Hoelzel, A. R., & Dover, G. A. (1991). Genetic differentiation between sympatric killer whale populations. Heredity, 66(2), 191.

Hoelzel, A. R., Hey, J., Dahlheim, M. E., Nicholson, C., Burkanov, V., & Black, N. (2007). Evolution of population structure in a highly social top predator, the killer whale. Molecular Biology and Evolution, 24(6), 1407–1415.

Holmes I, Davis Rabosky AR. (2018) Natural history bycatch: a pipeline for identifying metagenomic sequences in RADseq data. PeerJ 6:e4662 https://doi.org/10.7717/peerj.4662

Huerta-Cepas, J., Forslund, K., Coelho, L. P., Szklarczyk, D., Jensen, L. J., von Mering, C., & Bork, P. (2017). Fast genome-wide functional annotation through orthology assignment by eggNOG-mapper. Molecular biology and evolution, 34(8), 2115–2122.

Huson, D. H., Auch, A. F., Qi, J., & Schuster, S. C. (2007). MEGAN analysis of metagenomic data. Genome research, 17(3), 377–386.

Hyatt, D., Chen, G. L., LoCascio, P. F., Land, M. L., Larimer, F. W., & Hauser, L. J. (2010). Prodigal: prokaryotic gene recognition and translation initiation site identification. BMC bioinformatics, 11(1), 119.

Johnson, S. S., Zaikova, E., Goerlitz, D. S., Bai, Y., & Tighe, S. W. (2017). Real-time DNA sequencing in the Antarctic dry valleys using the Oxford Nanopore sequencer. Journal of Biomolecular Techniques: JBT, 28(1), 2.

Jones, F. C., Grabherr, M. G., Chan, Y. F., Russell, P., Mauceli, E., Johnson, J., Swofford, R., Pirun, M., Zody, M.C., White, S. et al. (2012). The genomic basis of adaptive evolution in threespine sticklebacks. Nature, 484(7392), 55.

Kircher, M., Sawyer, S., & Meyer, M. (2011). Double indexing overcomes inaccuracies in multiplex sequencing on the Illumina platform. Nucleic acids research, 40(1), e3–e3.

Kolodny, O., Weinberg, M., Reshef, L., Harten, L., Hefetz, A., Gophna, U., Feldman, M. W., & Yovel, Y. (2017). Who is the host of the host-associated microbiome? Colony-level dynamics overshadow individual-level characteristics in the fur microbiome of a social mammal, the Egyptian fruit-bat. bioRxiv, 232934.

Kong, H. H., Oh, J., Deming, C., Conlan, S., Grice, E. A., Beatson, M. A., Nomicos, E., Polley, E.C., Komarow, H.D., Murray, P.R. & Turner, M. L. (2012). Temporal shifts in the skin microbiome associated with disease flares and treatment in children with atopic dermatitis. Genome research, 22(5), 850–859.

Kong, H. H., & Segre, J. A. (2012). Skin microbiome: looking back to move forward. Journal of Investigative Dermatology, 132(3), 933–939.

Konishi, K., Tamura, T., Zenitani, R., Bando, T., Kato, H., & Walløe, L. (2008). Decline in energy storage in the Antarctic minke whale (Balaenoptera bonaerensis) in the Southern Ocean. Polar Biology, 31(12), 1509–1520.

Koskella, B., Hall, L. J., & Metcalf, C. J. E. (2017). The microbiome beyond the horizon of ecological and evolutionary theory. Nature ecology & evolution, 1(11), 1606.

Kueneman, J. G., Parfrey, L. W., Woodhams, D. C., Archer, H. M., Knight, R., & McKenzie, V. J. (2014). The amphibian skin-associated microbiome across species, space and life history stages. Molecular Ecology, 23(6), 1238–1250.

Lax, S., Smith, D. P., Hampton-Marcell, J., Owens, S. M., Handley, K. M., Scott, N. M., Gibbons, S.M., Larsen, P., Shogan, B.D., Weiss, S., & Metcalf, J. L. (2014). Longitudinal analysis of microbial interaction between humans and the indoor environment. Science, 345(6200), 1048–1052.

Lemieux-Labonté, V., Tromas, N., Shapiro, B. J., & Lapointe, F. J. (2016). Environment and host species shape the skin microbiome of captive neotropical bats. PeerJ, 4, e2430.

Ley, R. E., Lozupone, C. A., Hamady, M., Knight, R., & Gordon, J. I. (2008). Worlds within worlds: evolution of the vertebrate gut microbiota. Nature Reviews Microbiology, 6(10), 776.

Leyden, J. J., McGiley, K. J., Mills, O. H., & Kligman, A. M. (1975). Age-related changes in the resident bacterial flora of the human face. Journal of Investigative Dermatology, 65(4), 379–381.

Li, H. (2013). Aligning sequence reads, clone sequences and assembly contigs with BWA-MEM. arXiv preprint arXiv:1303.3997.

Li, H., & Durbin, R. (2009). Fast and accurate short read alignment with Burrows-Wheeler transform. Bioinformatics, 25(14), 1754–1760.

Li, H., Handsaker, B., Wysoker, A., Fennell, T., Ruan, J., Homer, N., … & Durbin, R. (2009). The sequence alignment/map format and SAMtools. Bioinformatics, 25(16), 2078–2079.

Li, H., Qu, J., Li, T., Li, J., Lin, Q., & Li, X. (2016). Pika population density is associated with the composition and diversity of gut microbiota. Frontiers in microbiology, 7, 758.

Lusk, R. W. (2014). Diverse and widespread contamination evident in the unmapped depths of high throughput sequencing data. PloS one, 9(10), e110808.

Mangul S., M Olde Loohuis L., Ori A., Jospin G., Koslicki D., Yang H. T., Wu T., Boks M. P., Lomen-Hoerth C., Wiedau-Pazos M, … & Ophoff, R. A. (2016) Total RNA Sequencing reveals microbial communities in human blood and disease specific effects. bioRxiv. 057570

Matkin, C. O., Barrett-Lennard, L. G., Yurk, H., Ellifrit, D., & Trites, A. W. (2007). Ecotypic variation and predatory behavior among killer whales (*Orcinus orca*) off the eastern Aleutian Islands, Alaska. Fishery Bulletin, 105(1), 74–88.

McFall-Ngai, M. J., Henderson, B., & Ruby, E. G. (Eds.). (2005). The influence of cooperative bacteria on animal host biology (Vol. 9). Cambridge University Press.

McKenzie, V. J., Bowers, R. M., Fierer, N., Knight, R., & Lauber, C. L. (2012). Co-habiting amphibian species harbor unique skin bacterial communities in wild populations. The ISME journal, 6(3), 588.

Li, D., Liu, C. M., Luo, R., Sadakane, K., & Lam, T. W. (2015). MEGAHIT: an ultra-fast single-node solution for large and complex metagenomics assembly via succinct de Bruijn graph. Bioinformatics, 31(10), 1674–1676.

Martinez Arbizu, P. (2017). pairwiseAdonis: Pairwise multilevel comparison using adonis. R package version 0.0.1.

Menke, S., Melzheimer, J., Thalwitzer, S., Heinrich, S., Wachter, B., & Sommer, S. (2017). Gut microbiomes of free-ranging and captive Namibian cheetahs: diversity, putative functions, and occurrence of potential pathogens. Molecular ecology.

Mollerup, S., Friis-Nielsen, J., Vinner, L., Hansen, T. A., Richter, S. R., Fridholm, H., … & Mourier, T. (2016). Propionibacterium acnes: disease-causing agent or common contaminant? Detection in diverse patient samples by next-generation sequencing. Journal of clinical microbiology, 54(4), 980–987.

Morin, P. A., Archer, F. I., Foote, A. D., Vilstrup, J., Allen, E. E., Wade, P., … & Bouffard, P. (2010). Complete mitochondrial genome phylogeographic analysis of killer whales (Orcinus orca) indicates multiple species. Genome research, 20(7), 908–916.

Morueta-Holme, N., Blonder, B., Sandel, B., McGill, B. J., Peet, R. K., Ott, J. E., … & Svenning, J. C. (2016). A network approach for inferring species associations from co-occurrence data. Ecography, 39(12), 1139–1150.

Moura, A. E., Janse van Rensburg, C., Pilot, M., Tehrani, A., Best, P. B., Thornton, M., … & Dahlheim, M. E. (2014). Killer whale nuclear genome and mtDNA reveal widespread population bottleneck during the last glacial maximum. Molecular biology and evolution, 31(5), 1121–1131.

Mukherjee, S., Huntemann, M., Ivanova, N., Kyrpides, N. C., & Pati, A. (2015). Large-scale contamination of microbial isolate genomes by Illumina PhiX control. Standards in genomic sciences, 10(1), 18.

Naccache, S. N., Greninger, A. L., Lee, D., Coffey, L. L., Phan, T., Rein-Weston, A., … & Chiu, C. Y. (2013). The perils of pathogen discovery: origin of a novel parvovirus-like hybrid genome traced to nucleic acid extraction spin columns. Journal of virology, 87(22), 11966–11977.

Nater, A., Mattle-Greminger, M. P., Nurcahyo, A., Nowak, M. G., de Manuel, M., Desai, T., … & Lameira, A. R. (2017). Morphometric, behavioral, and genomic evidence for a new Orangutan species. Current Biology, 27(22), 3487–3498.

Nelson, T. M., Apprill, A., Mann, J., Rogers, T. L., & Brown, M. V. (2015). The marine mammal microbiome: current knowledge and future directions. Microbiology Australia, 36(1), 8–13.

Oh, J., Byrd, A. L., Deming, C., Conlan, S., Barnabas, B., Blakesley, R., … Segre, J. A. (2014). Biogeography and individuality shape function in the human skin metagenome. Nature, 514(7520), 59–64. https://doi.org/10.1038/nature13786

Oksanen, J., Guillaume Blanchet, F., Kindt, R., & Legendre, P. (2017). others. 2016. vegan: Community ecology package. R package version 2.3–5.

Overbeek, R., Begley, T., Butler, R. M., Choudhuri, J. V., Chuang, H. Y., Cohoon, M., … & Fonstein, M. (2005). The subsystems approach to genome annotation and its use in the project to annotate 1000 genomes. Nucleic acids research, 33(17), 5691–5702.

Palsbøll, P. J., F. Larsen, And E. S. Hansen. 1991. Sampling of skin biopsies from free-ranging large cetaceans in West Greenland; Development of new biopsy tips and bolt designs. Rep. Int. Whaling Comm., Special Issue, 13: 71–79.

Parker, J., Helmstetter, A. J., Devey, D., Wilkinson, T., & Papadopulos, A. S. (2017). Field-based species identification of closely-related plants using real-time nanopore sequencing. Scientific reports, 7(1), 8345.

Parsons, K. M., Durban, J. W., Burdin, A. M., Burkanov, V. N., Pitman, R. L., Barlow, J., … & Wade, P. R. (2013). Geographic patterns of genetic differentiation among killer whales in the northern North Pacific. Journal of Heredity, 104(6), 737–754.

Paulson, J. N., Stine, O. C., Bravo, H. C., & Pop, M. (2013). Differential abundance analysis for microbial marker-gene surveys. Nature methods, 10(12), 1200.

Petersen, T. N., Lukjancenko, O., Thomsen, M. C. F., Sperotto, M. M., Lund, O., Aarestrup, F. M., & Sicheritz-Pontén, T. (2017). MGmapper: Reference based mapping and taxonomy annotation of metagenomics sequence reads. PloS one, 12(5), e0176469.

Phillips, C. D., Phelan, G., Dowd, S. E., McDonough, M. M., Ferguson, A. W., Delton Hanson, J., Siles, L., Ordóñez-Garza, N.I.C.T.É., San Francisco, M. & Baker, R. J. (2012). Microbiome analysis among bats describes influences of host phylogeny, life history, physiology and geography. Molecular ecology, 21(11), 2617–2627.

Piñeiro-Vidal, M., Gijón, D., Zarza, C., & Santos, Y. (2012). Tenacibaculum dicentrarchi sp. nov., a marine bacterium of the family Flavobacteriaceae isolated from European sea bass. International journal of systematic and evolutionary microbiology, 62(2), 425–429.

Pitman, R. L., & Durban, J. W. (2010). Killer whale predation on penguins in Antarctica. Polar Biology, 33(11), 1589–1594.

Pitman, R. L., & Durban, J. W. (2012). Cooperative hunting behavior, prey selectivity and prey handling by pack ice killer whales (Orcinus orca), type B, in Antarctic Peninsula waters. Marine Mammal Science, 28(1), 16–36.

Pitman, R. L., & Ensor, P. (2003). Three forms of killer whales (Orcinus orca) in Antarctic waters. Journal of Cetacean Research and Management, 5(2), 131–140.

Pitman, R. L., Fearnbach, H. & Durban, J. W. (2018). Abundance and population status of Ross Sea killer whales (Orcinus orca, type C) in McMurdo Sound, Antarctica: Evidence for impact by commercial fishing in the Ross Sea? Polar Biology DOI:/10.1007/s00300-017-2239-4

Poelstra, J. W., Vijay, N., Bossu, C. M., Lantz, H., Ryll, B., Müller, I., … & Wolf, J. B. (2014). The genomic landscape underlying phenotypic integrity in the face of gene flow in crows. Science, 344(6190), 1410–1414.

Quast C, Pruesse E, Yilmaz P, Gerken J, Schweer T, Yarza P, Peplies J, Glöckner FO (2013) The SILVA ribosomal RNA gene database project: improved data processing and web-based tools. Opens external link in new windowNucl. Acids Res. 41 (D1): D590–D596.

Quince, C., Walker, A. W., Simpson, J. T., Loman, N. J., & Segata, N. (2017). Shotgun metagenomics, from sampling to analysis. Nature biotechnology, 35(9), 833.

Quinlan, A. R., & Hall, I. M. (2010). BEDTools: a flexible suite of utilities for comparing genomic features. Bioinformatics, 26(6), 841–842.

Ranjan, R., Rani, A., Metwally, A., McGee, H. S., & Perkins, D. L. (2016). Analysis of the microbiome: advantages of whole genome shotgun versus 16S amplicon sequencing. Biochemical and biophysical research communications, 469(4), 967–977.

Rohland, N., & Reich, D. (2012). Cost-effective, high-throughput DNA sequencing libraries for multiplexed target capture. Genome research, 22(5), 939–946.

Romano-Bertrand, S., Licznar-Fajardo, P., Parer, S., & Jumas-Bilak, E. (2015). Impact de l’environnement sur les microbiotes: focus sur l’hospitalisation et les microbiotes cutanés et chirurgicaux. Revue Francophone des Laboratoires, 2015(469), 75–82.

Rothschild, D., Weissbrod, O., Barkan, E., Korem, T., Zeevi, D., Costea, P. I., … & Pevsner-Fischer, M. (2018). Environmental factors dominate over host genetics in shaping human gut microbiota composition. Nature doi:10.1038/nature25973

Salazar, G., & Sunagawa, S. (2017). Marine microbial diversity. Molecular Ecology, 27, R431–R510.

Salter, S. J., Cox, M. J., Turek, E. M., Calus, S. T., Cookson, W. O., Moffatt, M. F., … & Walker, A. W. (2014). Reagent and laboratory contamination can critically impact sequence-based microbiome analyses. BMC biology, 12(1), 87.

Der Sarkissian, C., Ermini, L., Schubert, M., Yang, M. A., Librado, P., Fumagalli, M., … & Petersen, B. (2015). Evolutionary genomics and conservation of the endangered Przewalski’s horse. Current Biology, 25(19), 2577–2583.

Saulitis, E., Matkin, C., Barrett-Lennard, L., Heise, K., & Ellis, G. (2000). Foraging strategies of sympatric killer whale (Orcinus orca) populations in Prince William Sound, Alaska. Marine mammal science, 16(1), 94–109.

Scharschmidt, T. C., & Fischbach, M. A. (2013). What lives on our skin: ecology, genomics and therapeutic opportunities of the skin microbiome. Drug Discovery Today: Disease Mechanisms, 10(3-4), e83–e89.

Schmieder, R., & Edwards, R. (2011). Quality control and preprocessing of metagenomic datasets. Bioinformatics, 27(6), 863–864.

Schubert, M., Lindgreen, S., & Orlando, L. (2016). AdapterRemoval v2: rapid adapter trimming, identification, and read merging. BMC research notes, 9(1), 88.

Segata, N., Izard, J., Waldron, L., Gevers, D., Miropolsky, L., Garrett, W. S., & Huttenhower, C. (2011). Metagenomic biomarker discovery and explanation. Genome biology, 12(6), R60.

Shotts Jr, E.B., Albert, T. F., Wooley, R. E., & Brown, J. (1990). Microflora associated with the skin of the bowhead whale *(Balaena mysticetus*). Journal of Wildlife Disease, 26, 351–359.

Song, S. J., Lauber, C., Costello, E. K., Lozupone, C. A., Humphrey, G., Berg-Lyons, D., Caporaso, J. G., Knights, D., Clemente, J. C., Nakielny, S. & Gordon, J. I. (2013). Cohabiting family members share microbiota with one another and with their dogs. elife, 2.

Sunagawa, S., Coelho, L.P., Chaffron, S., Kultima, J.R., Labadie, K., Salazar, G., Djahanschiri, B., Zeller, G., Mende, D.R., Alberti, A., et al. (2015). Structure and function of the global ocean microbiome. Science 348, 1261359.

Sun, C., Fu, G. Y., Zhang, C. Y., Hu, J., Xu, L., Wang, R. J., … & Zhang, X. Q. (2016). Isolation and complete genome sequence of Algibacter alginolytica sp. nov., a novel seaweed-degrading Bacteroidetes bacterium with diverse putative polysaccharide utilization loci. Applied and environmental microbiology, 82(10), 2975–2987.

Tessler, M., Neumann, J. S., Afshinnekoo, E., Pineda, M., Hersch, R., Velho, L. F. M., … & Mason, C. E. (2017). Large-scale differences in microbial biodiversity discovery between 16S amplicon and shotgun sequencing. Scientific reports, 7(1), 6589.

Tung, J., Barreiro, L. B., Burns, M. B., Grenier, J. C., Lynch, J., Grieneisen, L. E., … & Archie, E. A. (2015). Social networks predict gut microbiome composition in wild baboons. Elife, 4.

Võgene, õ. J., Herbig, A., Campana, M. G., García, N. M. R., Warinner, C., Sabin, S., … & Bos, K. I. (2018). Salmonella enterica genomes from victims of a major sixteenth-century epidemic in Mexico. Nature Ecology & Evolution, 2, 520.

van der Valk T, Vezzi F, Ormestad M, Dalén L, Guschanski K. Low rate of index hopping on the Illumina HiSeq X platform. Biotechniques. BioRxiv, doi: 10.1101/179028).

Walke, J. B., Becker, M. H., Loftus, S. C., House, L. L., Cormier, G., Jensen, R. V., & Belden, L. K. (2014). Amphibian skin may select for rare environmental microbes. The ISME journal, 8(11), 2207.

Warinner, C., Herbig, A., Mann, A., Fellows Yates, J. A., Weiß, C. L., Burbano, H. A., … & Krause, J. (2017). A robust framework for microbial archaeology. Annual review of genomics and human genetics, 18, 321–356.

Wright, E. S., & Vetsigian, K. H. (2016). Inhibitory interactions promote frequent bistability among competing bacteria. Nature communications, 7, 11274.

Wright, E. S., & Vetsigian, K. H. (2016). Quality filtering of Illumina index reads mitigates sample cross-talk. BMC genomics, 17(1), 876.

Wolz, M., Carly, R., Yarwood, S. A., Grant, E. H. C., Fleischer, R. C., & Lips, K. R. (2017). Effects of host species and environment on the skin microbiome of Plethodontid salamanders. Journal of Animal Ecology.

Ying, S., Zeng, D. N., Chi, L., Tan, Y., Galzote, C., Cardona, C., Lax, S., Gilbert, J. & Quan, Z. X. (2015). The influence of age and gender on skin-associated microbial communities in urban and rural human populations. PloS one, 10(10), e0141842.

Zhang C., Cleveland K., Schnoll-Sussman F., McClure B., Bigg M., Thakkar P., Schultz N., Shah M. A., Betel D (2015) Identification of low abundance microbiome in clinical samples using whole genome sequencing. Genome Biology, 16(1).

